# An altered metabolism in leukocytes showing in vitro igG memory from SARS-CoV-2-infected patients

**DOI:** 10.1101/2021.05.27.445918

**Authors:** G. Fanelli, F. Gevi, G. Zarletti, M. Tiberi, V. De Molfetta, G. Scapigliati, A.M. Timperio

## Abstract

Coronavirus disease 2019 (COVID 19) is a systemic infection that exerts a significant impact on cell metabolism. In this study we performed metabolomic profiling coupled with multivariate statistics analysis obtained from 43 in vitro cultures of peripheral blood mononuclear cells (PBMC), 19 of which displaying IgG memory for spike-S1 antigen 60-90 days after infection. By using mass spectrometry analysis, a significant up regulation of S-adenosyl-Homocysteine, Sarcosine and Arginine was found in leukocytes showing IgG memory. These metabolites are known to be involved in physiological recovering from viral infections and immune activities, and our findings might represent a novel and easy measure that could be of help in understanding SARS-Cov-2 effects on leukocytes.

## Introduction

The SARS-CoV-2 virus induces an unprecedented pandemic (COVID-19) characterized by a range of respiratory symptoms that may progress to acute respiratory distress syndrome (ARDS), and multi-organ dysfunction (1). SARS-CoV-2 triggers both innate and specific immune responses, and once the virus gains access within the target cell, the host’s immune system mainly recognizes its surface epitopes including the spike-S1 protein (2). Among other effects, a recent study showed that COVID-19 leads to a reduced population of regulatory lymphocytes, inducing an increase in inflammatory responses, in cytokines production, and proceeds toward tissue damage and systemic deficiency of organs (3). It is conceivable to speculate that at later stages after infection, patients that recovered from the pathology developed an immune response with memory and maintain antiviral defence, but the cellular physiological pathways underlying these mechanisms remain to be better understood. To this, a knowledge on cellular processes that could be related to infection can be achieved through metabolomics, an emerging tool applied to systems immunology platforms and already employed in COVID-19 studies (4). The aim of this preliminary study was to investigate possible relationships between leukocytes displaying an IgG antibody memory to SARS-Cov-2 (5) and their metabolic profile, in order to detect possible metabolites involved in late stages of antiviral defences.

## Results

Metabolites were extracted from PBMC cultures displaying an anti spike-S1 IgG-memory (IgGm+; N=19) and from PBMC without measurable IgG-memory (IgGm-; N=24) measured by Cell-ELISA 60-90 days after SARS-Cov-2 screening. Datasets shows age/gender and Cell-ELISA levels for each subject. We use MetaboAnalyst 5.0 platforms to perform untargeted metabolomics and identify the most relevant metabolites altered in IgGm+ subjects, and results indicate that the two different groups are well clustered, as shown in Figure 1A by the supervised PLS-DA, with red spots representing IgGm+ samples and green spots IgGm-. Multivariate analysis suggests significant variations present in IgGm+ subjects. The loading plot of metabolites in the analyzed samples projected into the PLS-DA model is shown in Figure 1B. The significantly-discriminated metabolites were identified using the Volcano plot analysis (Fig 1C). The univariate analysis identified significant accumulation of specific metabolites most of which were expressed in IgGm+ leukocytes. The metabolites displaying a major alteration were sarcosine, s-adenosyl-homocysteine and arginine.

**Fig. 1.**
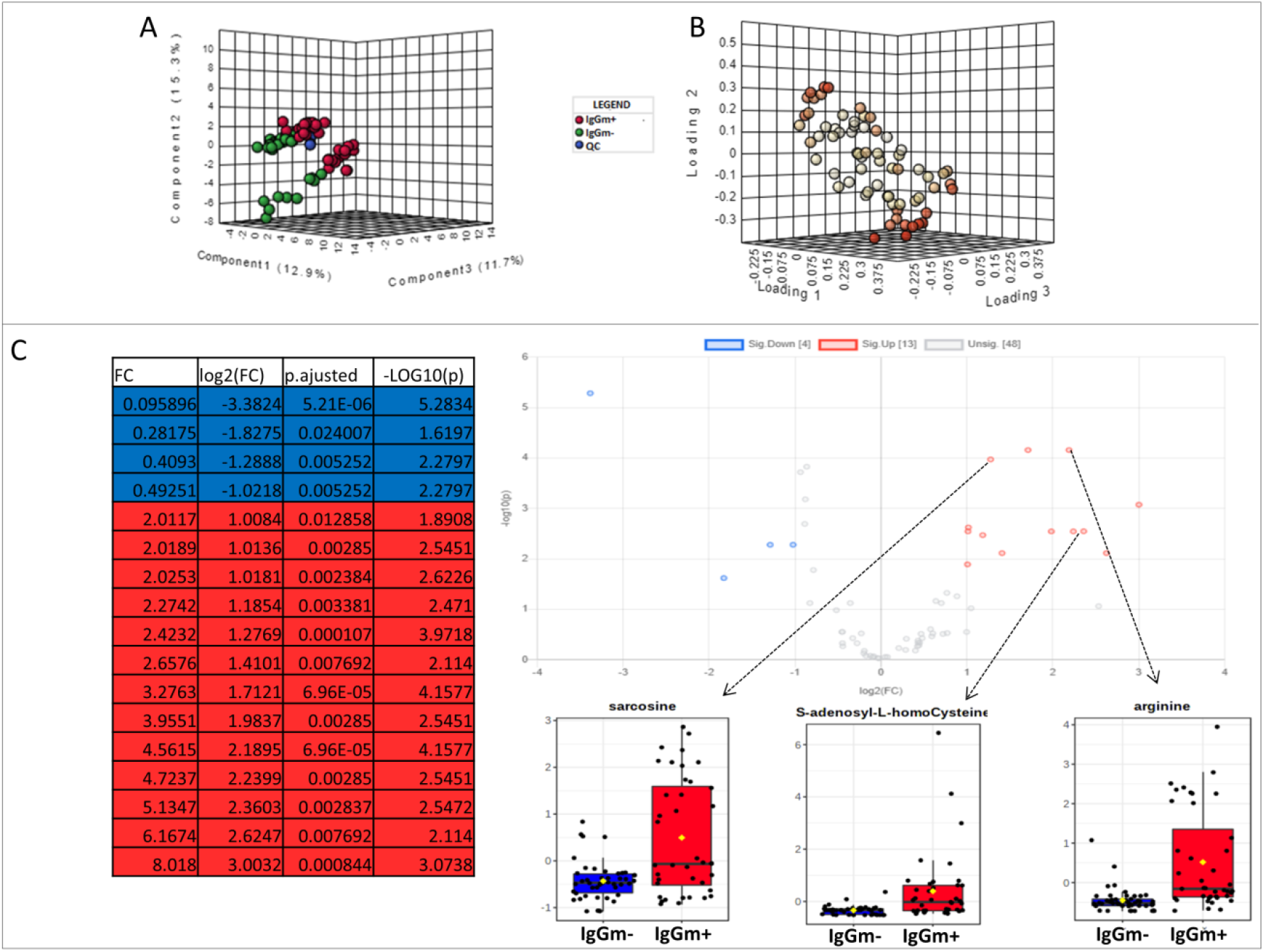
(A) Multivariate statistical analysis based on untargeted metabolite profile data derived from leukocytes of patients IgGm+ (Red), IgGm- (green) and QC (Blue). (B) shows the loading plot of the metabolites in the analyzed samples projected into the PLS-DA model. (C) Volcano plot showing the most significant metabolites found by univariate analysis. The levels of metabolites were significantly different in the IgGm+ compared to IgGm-. The volcano plot summarizes both fold-change and t-test criteria for all metabolites. The blue-highlighted table shows metabolites up-regulated in IgGm-leukocytes, whereas in the red-highlighted metabolites up-regulated in IgGm+ leukocytes. The more significantly altered metabolites in IgGm+ leukocytes are sarcosine, s-adenosyl-homocysteine, arginine

## Discussion

The SARS-Cov-2 induces an antibody memory against spike-S1 antigen that can be measured by in vitro assays in PBMC from two months onward after infection (5), and the aim of our work was to investigate metabolic features in these conditions. Through metabolomic analysis, the results showed that in vitro cultured and unstimulated PBMC display a significant up-regulation of metabolically-related metabolites sarcosine, s-adenosyl-L-homocysteine and arginine.

The S-adenosyl-methionine (AdoMet, also frequently abbreviated as SAM and SAMe) is widely known as a main biological methyl donor converted to s-adenosyl-L-homocysteine (AdoHcy), and Sarcosine as an acceptor of methyl group. AdoHcy cellular content is regulated by the enzyme AdoHcy hydrolase (AHCY), which reversibly cleaves this molecule into adenosine and homocysteine (Hcy). The thermodynamics of AHCY stimulates AdoHcy biosynthesis rather than hydrolysis, and in vivo the reaction proceeds in the direction of hydrolysis only if the products, adenosine and Hcy, are rapidly removed. This is essential to prevent accumulation of AdoHcy (fig2) which is a potent inhibitor of methylation reactions (6).

**Fig. 2.**
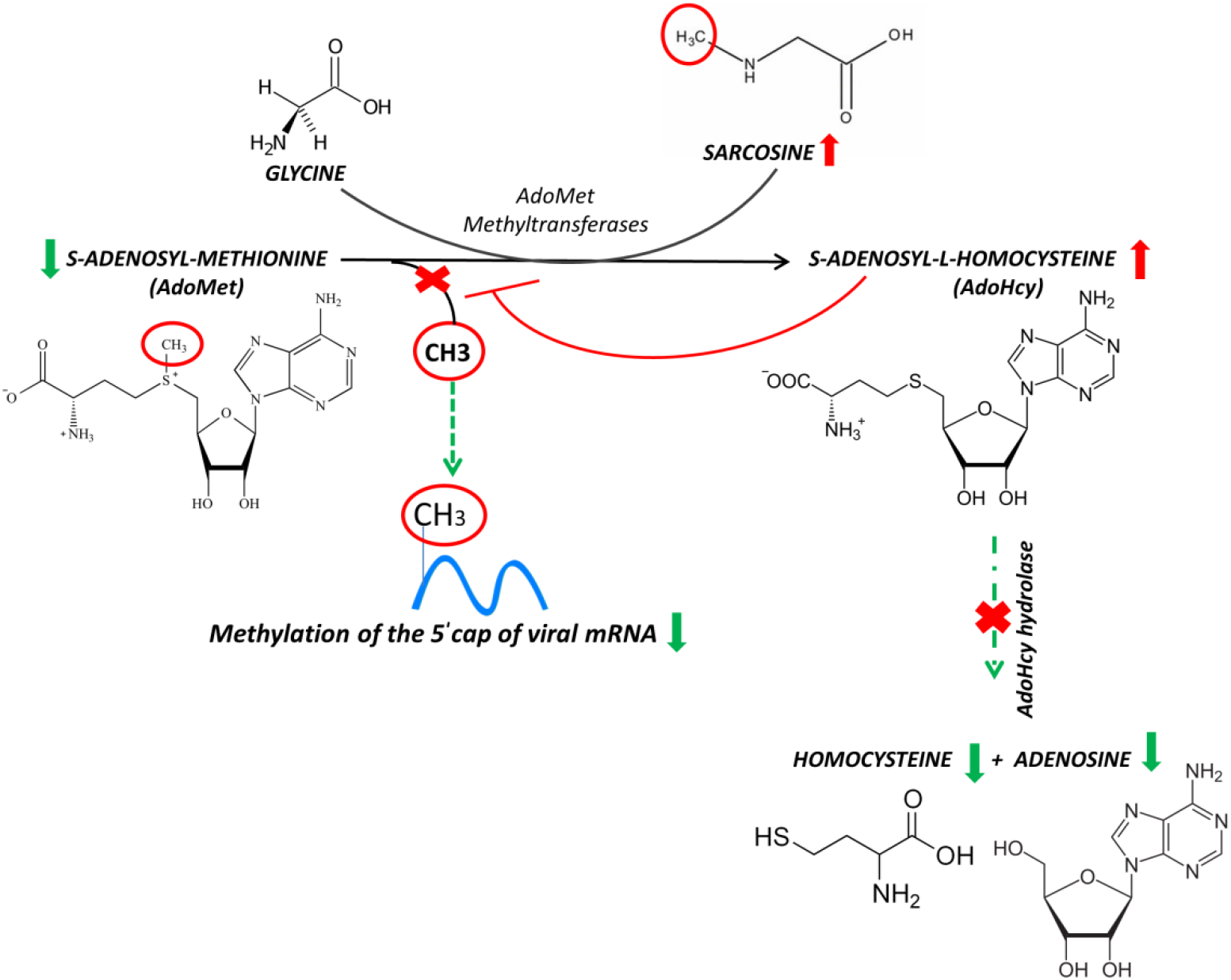
Reactions of methionine metabolism. S-adenosyl-methionine (SAM) donates its methyl group to acceptor molecules, sarcosine, generating S-adenosylhomocysteine (SAH) limiting methylation of the 5’ cap of viral messenger RNA.

The ratio of AdoMet to AdoHcy is frequently considered as a metabolic gauge controlling in vivo methylation reactions, where a decrease in this ratio predicts reduced methylation capacity (7). These methylations are required for the 5’-cap formation of viral mRNAs. The RNA cap has multiple roles in gene expression, including enhancement of RNA stability, splicing, nucleocytoplasmic transport, and translation initiation, necessary for viral RNA replication (8). In this respect, inhibition of the enzyme AdoHcy hydrolase can be used as a therapy against virus infection, because indirectly limits methylation of the 5’ cap of viral messenger RNA as already noticed for both Ebola virus and African swine fever virus (9,10). By our results, the strong increase of AdoHcy allowed us to hypothesize an inhibition of s-adenosyl-L-homocysteine hydrolase, and consequently leukocytes may proceed to virus elimination. Importantly, inhibition of this latter mechanism is a target in antiviral strategies employed to fight both African swine fever virus and Ebola virus. It is also conceivable to conclude that blocking capping viral mRNAs could be a potential target for antiviral therapies, due to the translation of viral proteins.

Interestingly, we also observed a strong up regulation of arginine in IgGm+ leukocytes, in addition to AdoHcy and sarcosine. Arginine is a key amino acid nutrient shown to be essential to the life cycle of many viruses, although having a controversial role in SARS-CoV-2 infection. According to a previous work (11), the plasma of COVID 19 patients contains a lower amount of arginine and, as a consequence, its depletion could potentially delay and/or compromise intensive care unit recovery (11). Nevertheless, arginine was shown to increase the viral replication in SARS-CoV-2 infected cells. For instance, the nucleocapsid (N) protein contains 6.9 % of arginine which interacts with the negatively charged RNA strands, and helps to bind and wrap to have efficient virion packing. Additionally, arginine residues on the spike S1 protein may be crucial to stabilize viral interaction with the host cell receptor angiotensin-converting enzyme 2 to facilitate viral entry (12).With respect to immune defences, it should be noted that an increase in arginine concentration may play a role in the regulation of immune cell reactivity through the proliferation and differentiation of naive T cells to memory (13). Arginine could preferentially enhance the proliferation of T lymphocyte subpopulations by increasing specific receptor expression and IL-2 production (14), and increase T cell number/responses (15). Moreover, an arginine increase could also enhance lymphocyte proliferation by providing a substrate for protein synthesis and/or precursors of polyamines or nitric oxide, being important in sustaining cellular proliferation (13). Together, our preliminary results suggest that unstimulated PBMC analyzed up to 90 days after SARS-CoV-2 infection show an increase in antiviral metabolites (sarcosine and S adenosylhomocysteine), and a modulation of arginine metabolism, involved in innate and adaptive immunity. With further work, the analysis of these metabolites might represent a biomarker of effective and long-standing antiviral activation of PBMC.

## Materials and Methods

43 subjects (24 males and 19 females, dataset) undergoing COVID-19 serological analysis (Centro Polispecialistico Giovanni Paolo I, Viterbo, I) were enrolled in this study from October 2020 to March 2021. All participants provided informed written consent to participate in the research project, and the study was approved by the Regional Ethical Board in Ospedale L. Spallanzani, Roma, (number 169, approval 22/07/2020) and, in accordance with the Helsinki Declaration, written informed consent was obtained from all subjects. Determination of in vitro IgG B cell memory for spike-S1 virus protein and of specific IgG were performed by Cell-ELISA in PBMC, from all donors as previously described without modifications (5), employing spike-S1 coated wells from a commercial source, (details in datasets), experimental outputs were net absorbance values (A450 nm) with background subtracted. Parallel cultures of PBMC employed in Cell-ELISA were incubated without stimulants at 106/ml at 37°C for 48 h in 100 ul/well of RPMI medium containing 10% FCS and antibiotics, then centrifuged at 500xg, and pelleted cells immediately employed for metabolomic analysis. Metabolites were extracted and analysed by LC-MS following the protocol published in (16). 10 μL aliquots of all samples were pooled as a quality control (QC) sample which was run every 10 samples. Replicates were exported as.mzXML files and processed through MAVEN.5.2. Multivariate (PLS-DA) and Univariate (Volcano plot) statistical analyses were performed on the entire metabolomics data set using the MetaboAnalyst 5.0 software.

## Author Contributions

Samples were collected by Z.G. T.A.M and S.G conceived and designed the experiment; F.G and G.F performed the experiments and contributed analysis tools; T.A.M, F.G and G.F wrote the paper with the input of S.G. T.A.M., F.G, G.F, S.G, Z.G, T.M and D.M.V reviewed and edited the manuscript. All authors have read and agreed to the published version of the manuscript.

## Acknowledgments

This work was supported by PRIN 2017 from University of Viterbo. and by Ministero dell’Istruzione, dell’Università e della Ricerca (MIUR), Rome, Italy.

## dataset

age/gender and In vitro IgG memory. *Cell-ELISA absorbance (A450 nm) net values, positive values are in bold with a cutoff value of 0.07. The Cell-ELISA protocol and cutoff value are as previously described (5).

